# In-vivo Measurement of Radio Frequency Electric Fields in Mice Brain

**DOI:** 10.1101/2022.08.16.504138

**Authors:** Omid Yaghmazadeh, Seth Schoenhardt, Arya Sarabandi, Ali Sabet, Kazem Sabet, Fatemeh Safari, Leeor Alon, György Buzsáki

**Author notes:** Correspondence: György Buzsáki (Neuroscience Institute, NYU School of Medicine, 435 E30th St, New York University, New York, NY 10016, USA), Omid Yaghmazadeh (Neuroscience Institute, NYU School of Medicine, 435 E30th St, New York, NY 10016, USA), and Leeor Alon (Department of Radiology, NYU School of Medicine, 660 1^st^ Ave, New York, NY 10016, USA).

## Abstract

With development of novel technologies, radio frequency (RF) energy exposure is expanding at various wavelengths and power levels. These developments necessitate updated approaches of RF measurements in complex environments, particularly in live biological tissue. In this study, we introduce a technique for direct in-vivo measurement of electric fields in living tissue. Proof of principle in-vivo electric field measurements were conducted in rodent brains using Bismuth Silicon Oxide (BSO) crystals exposed to varying levels of RF energy. Electric field measurements were calibrated and verified using in-vivo temperature measurements using optical temperature fibers alongside electromagnetic field simulations of a transverse electromagnetic (TEM) cell.

**SIGNIFICANCE:** Accurate dosimetry of the absorbed radio frequency (RF) electric fields (E-Fields) by the live tissue is the keystone of environmental health considerations for this type of ever-growing non-ionizing radiation energy. The complexity of biological tissue and technical difficulties have made direct measurement of E-fields in live tissue challenging leading to application of ex-vivo and in-silico approaches. Here, we present a novel method for in-vivo direct measurement of RF E-fields in anesthetized mice brain using electro-optic sensors.

## INTRODUCTION

Since the primary Radio Frequency (RF) applications in the early 20^th^ century, the interaction of RF energy with biological tissue and its potential risks have gained a great deal of interest (^1–4^). With advancement of novel technologies, the exposure to radio frequency (RF) energy has never stopped expanding. The extent of RF applications and the increase in applied frequency bands have resurrected questions with regards to RF energy exposure and its impact on humans (^5^). For example, development of high power microwave (HMP) applications for civil and military purposes has generated a new wave of concerns for health implication of RF energy radiation on the brain and the body (^6,7^). In line with such concerns, the International Commission on Non-Ionizing Radiation Protection (ICNIRP) has recently updated their guidelines for RF energy exposure limits (^8^).

Studying the effects of RF energy on biological tissue would require methods to accurately measure the absorbed RF energy in the tissue for which the widely used standard parameter is the specific absorption rate (SAR) (^8^). For example, assessment of the induced RF energy inside biological tissue is essential in various medical interventions (such as Magnetic Resonance Imaging, RF ablation, etc.). Furthermore, it is the core element for studying safety measures related to any RF energy exposure (such as cell phone usage, HPM applications, etc.). SAR and electric field (E-field) can be calculated either experimentally or by numerical simulations. Evidently, experimental assessments are preferable as numerical simulations are often vulnerable to errors in various ways. Due to the complexity of biological systems and the limitations of the E-field probes, it is difficult to conduct measurements in vivo, and as a result, RF compliance measurements are conducted in phantoms that simulate human tissue. However, those compliance measurements may not faithfully reflect real-life scenarios of exposure. Furthermore, many groups are using electromagnetic (EM) field simulations to assess exposure from various devices, but real-life complexities of the environment and tissue anatomy seldom can be replicated in-silico. Both EM field simulations and phantom compliance tests rely on tabulated dielectric properties of tissues measured in euthanized sheep (^9^). However, it is well known that, in live tissue, the physiological variation can greatly impact those dielectric properties (e.g., water content and hydration), thus reliance on these values may introduce non-negligible errors in RF exposure estimation.

From the materials point of view, the biological tissue constitutes very complex structures with variable composition across different subjects. Although their physical properties have been subject to extensive studies (^10^), providing several numerical or phantom representatives, their complexity requires direct experimental measurements for accurate examination. Therefore, different biological tissues should be considered as a stand-alone complex material and be examined accordingly especially when accurate measurements are demanded.

Experimental measurement of in-situ E-field in biological tissue (or phantom preparations with similar physical properties) induced by RF energy exposure can be performed either by temperature measurements (^11^) or using E-field probes (^12^). Using temperature measurements to assess the in situ E-field is reliable but only for RF energy exposure levels that induce a sufficient temperature rise in the tissue. E-field probes are usually constructed in two ways: metal based (^12^) or electro-optic (EO) probes (^13^). EO E-field probes have several advantages, such as their non-metallic nature (which makes them minimally invasive to the E-field distribution), wide bandwidth (from DC to several THz, ^14,15^), very high (pico-sec) temporal resolution (^16^), and relatively small dimensions, making them ideal for the in vivo measurement of the E-field. For example, recently developed HMP applications apply very short (µsec range) and very high power (KW/cm^2^ to MW/cm^2^) RF pulses (^6^). Measuring the in-situ E-field in biological tissue exposed to such radiation requires both high measuring range and high temporal resolution.

The possibility of direct measurement of induced RF E-fields inside the tissue (either *in-vivo* or *ex-vivo*) provides a more accurate assessment of the in-situ E-Field in living species exposed to such radiation. While E-field measurement in ex-vivo and phantom preparations provide a good insight on the amount of absorbed energy due to RF exposure, in-vivo measurements provide a direct and more accurate evaluation. In this study we introduce a methodology for direct measurement of RF E-fields in mice brain in-vivo where E-field measurements were calibrated and validated by temperature measurements conducted by optical temperature fibers alongside electromagnetic field simulations in a transverse electromagnetic (TEM) cell.

## RESULTS

We performed in situ E-field measurements in the brain of head-fixed anesthetized mice exposed to RF energy radiation using a BSO based E-field EO probe.

### Electro-Optical E-field measurement system

RF E-field measurements were performed by the commercially available NeoScan System manufactured by EMAG Technologies Inc. Fig. 1 shows the schematics and photo of the electro-optic field probe and the E-field measurement system. The E-field probe consists of an electro-optic crystal cut along one of its optical axes and mounted at the tip of an optical fiber via a graded index (GRIN) lens. The EO probe is mounted either on a fixed stand or on a computer-controlled translation stage and is then immersed in the E-field of the device under test (DUT; Fig. 1B). A continuous wave (CW) laser beam propagating through the optical fiber and then into the EO crystal experiences the Pockels effect, whereby the external E-field penetrating the EO crystal induces a change in its refractive index. As result, the polarization vector of the optical beam is rotated by an angle that is directly proportional to the strength of the E-field (^17^). The change in the polarization state in turn produces a measurable change in the optical intensity of the beam as it passes through a polarization analyzer. Finally, a high-speed photodetector at the end of the optical bench transduces the intensity of the input optical beam into a high-frequency electric signal. The output voltage of the photodetector is linearly proportional to the E-field of the device under test. Because of the extremely small values of Pockels coefficients, the output voltage of the photodetector is very weak and often buried under the system’s noise floor. An ultra-wideband multi-stage low-noise amplifier (LNA) is used to amplify this signal to a measurable level. In a frequency-domain near-field measurement configuration as shown in Fig. 1B, the amplified RF signal at the output of the photodetector is mixed with a local oscillator (LO) and is down-converted to an intermediate frequency (IF) of 100 MHz. Then a lock-in amplifier is utilized to measure both the amplitude and phase of this low-frequency signal, whose amplitude is directly proportional to the field strength of the device under test. A BSO probe with a <100> crystal cut can measure the normal component of the E-field of the DUT, while a BSO probe with a <110> crystal cut can measure the tangential E-field component.

**Figure 1.**
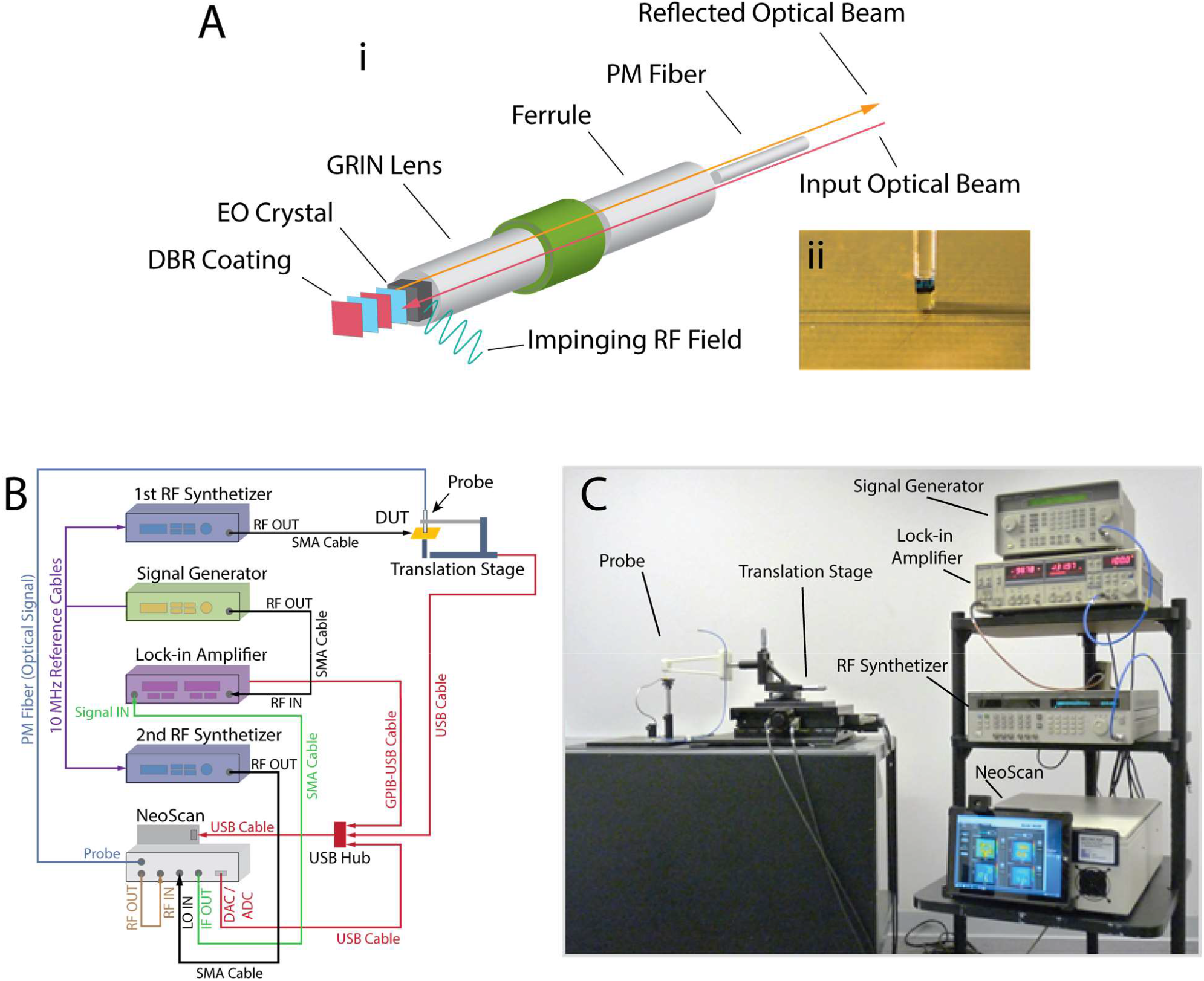
Electro-Optical E-field measurement system and its principal components. **A)** 3D drawing (i) and close-up photo (ii) of the ‘non-metallic’ electro-optic field probe. **B)** Schematic of the E-field measurement system. **C)** Photo of a NeoScan E-field measurement setup.

The EO E-field probe is made of all dielectric materials with no metallic components and can be brought to the very-near-field region of the DUT in a non-invasive manner. Due to its small footprint (1 mm^2^), it can be used to measure the E-fields with high spatial resolution with minimal distortion to incumbent EM waves. The minimum sampled space can be less than 10 μm corresponding to the focused laser beam spot size within the EO crystal. Another advantage of the EO probe is an extremely wide bandwidth (1 kHz – 40 GHz or higher). The probe can be calibrated to measure the absolute magnitude of E-fields over a wide dynamic range (0.1 V/m–1 MV/m).

### E-field probe calibration for measurement in biological tissue

In the measurement system for our EO probe, the output signal power, which is measured using either a spectrum analyzer or a lock-in amplifier, is proportional to the square of the E-field inside the BSO crystal:

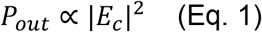

where *P*_*out*_ (W) and *E*_*c*_ (V.m^-1^) are the measured power by the system and the E-field inside the crystal, respectively. Therefore, for every power value measured by the system, the E-field inside the crystal is given by:

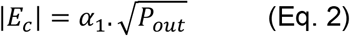

where *α*_1_ (V.m^-1^.W^-1/2^) is a constant defined as the crystal to output calibration factor. Its value depends on the electronic attenuation of the system, BSO dimension, and gain of the LNA. The value of the E-field inside the crystal depends not only on the incident E-field that the probe is exposed to, but also on the physical characteristics of the surrounding medium (or more precisely the interface between the medium and the sensor’s crystal). In other words, the E-field inside the crystal differs from the E-field inside the medium, in which it is placed in, due to the changes in dielectric properties at the interface of the medium material and the crystal. Therefore, the measurement system should be calibrated for the usage of each probe for application in a given medium (including air).

When the probe is placed inside a medium with a relatively homogenous distribution of the E-field (which is the case for the situation where the dimensions of the probe is smaller than the wavelength in the medium), the induced E-field inside the crystal is scaled to the E-field in the medium (Supp. Fig. 2C and 2D, ^18^) by:

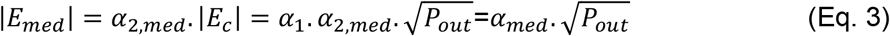

where *E*_*med*_ (V.m^-1^), *α*_2,*med*_ (a.u.) are the E-field in the medium surrounding the probe and a constant defined as the medium to crystal calibration factor, respectively. *α*_*med*_ = *α*_1_. *α*_2,*med*_ (V.m^-1^.W^-1/2^) is the total calibration factor, a medium-specific constant.

**Figure 2.**
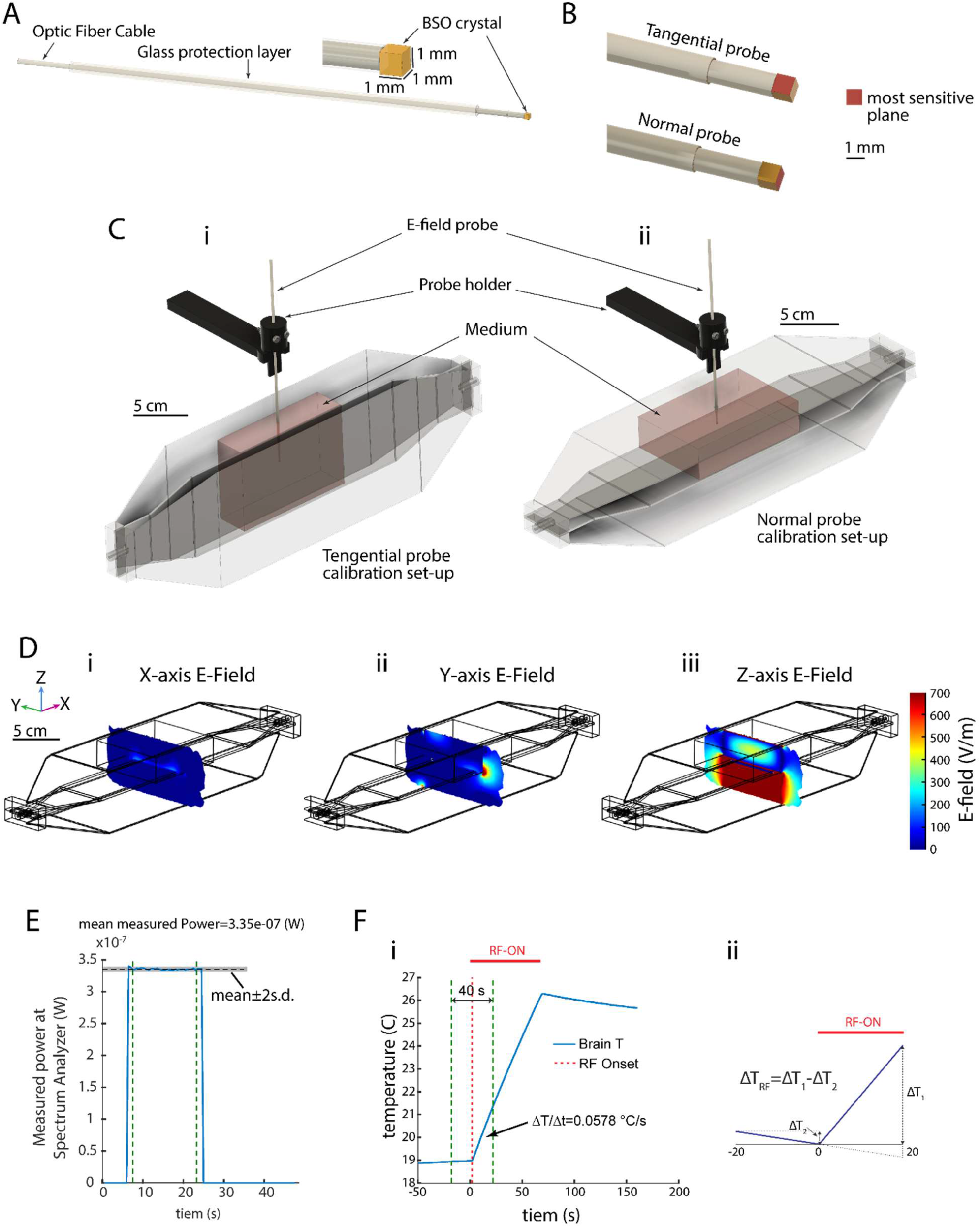
Calibration of electrooptic E-field probes for measurements in a medium. **A)** Schematic of the E-field probe used in these experiments with a cubic (1mm^3^) BSO crystal sensor. **B**) Schematic showing the sensitive plane on different probe types (tangential and normal). **C**) 3D schematic of the calibration set-up configuration for tangential (*left*) and normal (*right*) probes. E-field probe is placed in the center of a reservoir filled with the medium of interest (e.g., brain tissue) and placed at the middle of one of TEM cell’s chambers. **D**) Numerical simulations (COMSOL Metaphysics software) demonstrate that the E-field along the axis perpendicular to the TEM cell’s septum (E_z_) is an order of magnitude higher than in other directions (E_x_ and E_y_). **E**) Example measurement curve showing non-calibrated power from the spectrum analyzer. **F**) *i*: Temperature measurement using an optical temperature probe performed using the same set-up configuration as in C-*left. ii*: A simple plot showing how ΔT is calculated considering the slope of the temperature changes during the baseline.

Each probe needs to be calibrated separately (i.e. its *α*_*med*_ should be determined) for application in a given medium. EO probes in air were calibrated using a known E-field in an air background such as the field inside a Transverse Electro Magnetic (TEM) cell antenna when excited with known RF power. For calibration of the probe inside a medium other than air, experiments containing such medium, with known dielectric properties and the values of the true E-field is required. For this purpose, similar to previous reports (^18^), we used a reservoir of the medium of interest inside a TEM cell (Supp. Fig. 1). For a given input power into the TEM cell, we calculate the E-field in the center of the reservoir via temperature measurements. It should be noted that the reservoir dimensions were verified in simulations (Fig. 2) to be sufficiently large for ensuring a homogeneous E-field in the location of the probe (i.e., center of TEM cell).

The EO probes that we used in our experiments (shown in Fig. 2A) have a directional preference, *i*.*e*., they are sensitive to the *in-situ* field only from one direction and come in two formats: *normal probes* which are sensitive to the E-field parallel to the main axis of the probe, and *tangential probes* which are sensitive to the E-field at one direction in the plane perpendicular to the main axis of the probe (Fig. 2B). Because E-field in center of the cavities of a TEM cell is maximal in the direction perpendicular to the surface of its plates and minimal in other directions, two separate configurations of EO placement with regard to the TEM cell were used to calibrate different probe type (normal and tangential; Fig. 2C), leading to two calibration factors for the normal and tangential probes, respectively.

For the calibration method to work, an induced E-field in the medium, generated by the TEM cell needed to be dominated by a single E-field polarization (*i*.*e*. perpendicular to the TEM cell’s plates, Z in Fig. 2D), rather than in the other directions (X and Y in Fig. 2D). We validated this by numerical simulations as illustrated in Fig. 2D. As shown in this figure the E-field along Z direction is >10x higher than along the X- and Y-directions.

Fig. 2C shows the experimental set-up prepared for calibration of our normal and tangential EO probes. In order to calibrate our probes for E-field measurement in mouse brain, mice brains were extracted from healthy animals that were assigned to be sacrificed at the end of the experiments, collected from several laboratories in our institute. This prevented sacrificing the life of new animals for this purpose. Brain extraction was performed as close in time as possible to the experiment (one day before) and brain samples were kept in saline at 4°C. Right before the experiment, we removed the saline and made mixture of all brains. We filled the reservoir of our calibration set-up with this mixture and performed the probe calibration as discussed. Fig. 2E shows an example measurement (output power from the spectrum analyzer of the system) from a probe calibration experiment.

In order to calibrate the output of the system in each measurement one needs to verify the value of the actual E-field at the probe’s location (See Materials and Methods). Therefore, we conducted temperature measurements using an optical temperature probe (0.3mm diameter, PRB-329 100-01M-STM, Osensa Innovation Corp.) to measure the rate of the induced temperature increase and the corresponding Specific Absorption Rate (SAR) and consequently the E-field. We conducted these temperature measurements in the configuration for calibration of the tangential probe as shown in Fig. 2Ci. The result would apply to the calibration of the normal probe also as the temperature measurement is sensitive to the magnitude of the total E-field and not to its direction. Fig. 2Fi shows the change in temperature of the center of the brain mixture reservoir due to 70s of continuous-wave RF exposure (36W input power into the TEM cell). The resulting rate of T rise (ΔT/Δt) calculated by considering 20s before and after the onset of the RF exposure (Fig. 2Fi) is 0.0578 °C/s which leads to a calculated SAR and E-field values of 204.45 W/Kg and 617.64 V/m, respectively (see Materials and Methods, ^19^). During this short period after the onset of heating, heat removal from the system as well as heat diffusion across the brain is expected to be minimal, thus introducing small errors in the E field calculations. This statement is congruent with the linear temperature change observed in the brain at these heating time scales and intensities.

Dividing the un-calibrated measured power values (*P*_*out*_) obtained from the calibration experiments (1.13e-6 W and 3.28e-7 W for tangential and normal probe (average from three repetitions), respectively) by the actual E-field strength deducted from the temperature measurements the calibration factor (defined as 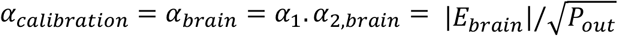 where *E*_brain_ (V.m^-1^) is the actual E-field value in the center of reservoir filled with the brain tissue) for the tangential and normal probes used in our experiments were 5.81e5 (V.m^-1^.W^-1/2^) and 1.08e6 (V.m^-1^.W^-1/2^), respectively. These factors will be used to adjust the E-field reading in the in vivo experiments (as shown in Fig. 3).

**Figure 3.**
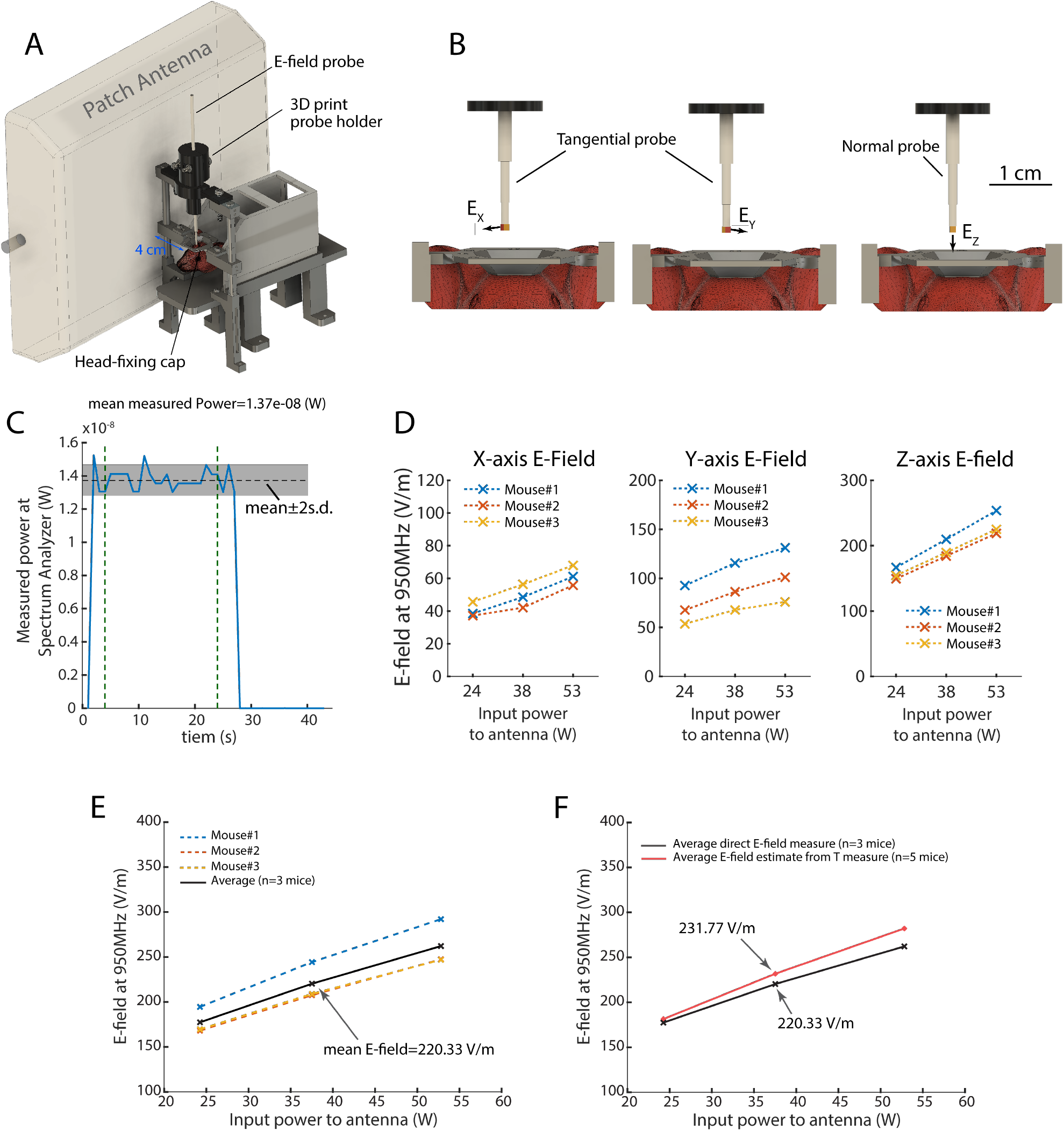
Direct measurement of E-field magnitude in mice brains *in-vivo*. **A)** Schematic of the experiment set-up: a mouse implanted with the EO probe is head-fixed and a patch antenna is placed next to the animal in a fixed position (4cm away from animal’s head). **B)** illustration of the measurement procedure to obtain E-fields along the x-,y-, and z-orientations. From left to right: first a tangential probe is implanted and E-field measurement is performed. Then the probe is extracted from the brain (but remains in the set-up) and turned 90° on the plane parallel to surface of the skull. The probe is implanted back in the brain and E-field is measured. Finally, the tangential probe is completely removed from the setup and a normal probe is implanted, and E-field is measured. **C)** Example curve showing non-calibrated power from the spectrum analyzer for in-vivo Y-axis measurement for 38W RF input power to antenna. **D)** Calibrated E-field magnitude values measured along the three axis shown in B (n=3 mice). **E)** Total E-field magnitude (n=3 mice) using data in D. **F)** Comparison between the E-field values obtained by direct measurement of E-field using EO probes (this study) and values estimated from in-vivo temperature measurements.

In order to establish a simpler way for calibrating our EO probes without the need to sacrifice a large number of animals, we examined our methodology on a phantom material with similar physical characteristics than the mice brain tissue. We developed a simple method to produce such phantom material from commercially available products (see Materials and Methods). We used a similar approach as that of probe calibration in brain tissue using temperature measurements. Supp. Fig. 3 shows the change in temperature of the center of the phantom material reservoir due to 70s of continuous-wave RF exposure (36W input power into the TEM cell). The resulting rate of T rise (ΔT/Δt) calculated by considering 20s before and after the onset of the RF exposure is 0.0373 °C/s which leads to a calculated SAR and E-field values of 168.15 W/Kg and 625.90 V/m, respectively (see Materials and Methods, ^19^). The power values (*P*_out_) measured in the calibration experiment with the phantom material were 9.80e-7 W and 2.67e-7 W for tangential and normal probe, respectively. As the phantom material we made has a high viscosity, it is possible to use temperature measurement to estimate the E-field. This is validated by comparing the calibration results with calibration coefficients of 6.32e5 (V.m^-1^.W^-1/2^) and 1.21e6 (V.m^-1^.W^-1/2^) for the tangential and the normal probes, respectively whose difference with the values obtained using the brain tissue sample are 8.8% and 12.3%, respectively.

### Direct measurement of RF E-field in mice brains in-vivo

We conducted experiments to measure *in-situ* E-field magnitude in the brain of urethane-anesthetized mice. We head-fixed the mice to ensure a similar RF E-field exposure for all experiment repetitions. We replicated the set-up from one of our recent studies where the effect of RF exposure on neuronal activity monitored by 1-photon Ca^2+^ imaging in-vivo was investigated (^20^).

Fig. 3A shows the experimental set-up for the head-fixed mouse with implanted EO probe in the brain. We measured the E-field values from three orthogonal directions using a normal and a tangential EO probe. As illustrated in Fig. 3B we first implanted the tangential EO probe and measured the E-field along the x axis (E_x_). Then, we extracted the tangential probe and rotated it for 90°, implanted it to the same depth in the brain and measured the E-field along the y axis (E_y_). Next, the tangential probe was extracted from the brain and replaced it with the normal probe which was then implanted to the same coordinates inside the brain and E-field along the z axis (E_z_) was recorded. Fig. 3D and 3E shows the results of the measurement of the magnitude of E-field along different directions (E_x_, E_y_, and E_z_) and the total E-field (E_tot_, see Materials and Methods). These results are calculated using the calibration coefficients obtained in probe calibration experiments in brain tissue.

As shown in Fig. 3F, the in-vivo E-field measurement results obtained in this study are in good agreement with the E-filed values driven from in-vivo temperature measurements (see Materials and Methods) performed by an optical temperature probe (1.1mm diameter, OTP-M, Opsens Solutions Inc.).

### Numerical simulations to assess the effects of material properties on the E-field measurement

We performed finite-element numerical simulations to study the effects of electrical conductivity (*σ*) and relative permittivity (*ε*_*r*_) of the medium on the induced E-field inside the crystal and consequently on the measurement. We created a 3D model of the experimental setup of the probe calibration procedure presented in Fig. 2C. We conducted simulation runs, using COMSOL Multiphysics (Comsol Inc.), with parametric sweeps for *σ*, varying its value from 0.4 to 4 (S.m^-1^) with 0.2 (S.m^-1^) incremental steps and for *ε*_*r*_, varying its value from 1 to 78.5 (a.u.) with 2.5 (a.u.) incremental steps. Supp. Fig. 2A shows the 3D design of the experiment in COMSOL. Supp. Fig. 2B illustrates the probe configuration used in the simulation including numerical probing volumes to measure the E-field at the center of the BSO crystal and at its adjacent volumes in the medium of interest. Supp. Fig. 2C and 2D confirm a linear relationship between the measured E-field (in any of the probing volumes) and the square root of the input power to the TEM cell and between any pair of the measured electric-field values for a given input power. As the EO probe output depends on the ratio between the E-field in the center of the crystal and the E-field at its neighboring medium (at its bottom for the normal probe and at one of its lateral sides for the tangential probe), we next examined this ratio as a function of the relative permittivity (Supp. Fig. 2E) and the electrical conductivity (Supp. Fig. 2F) of the medium for a fixed input voltage, hence a fixed input power (here 36 W similar to our probe calibration experiments), to the TEM cell and in the case of a normal probe as an example. Note the relatively low variability of the measured ratio between the E-field values when either the relative permittivity or the electrical conductivity of the medium is set close to those corresponding to the brain tissue (57 and 1 S/m, respectively) and varying the other. This shows that calibrating the EO probes using a phantom with a relative permittivity and an electrical conductivity close to the brain tissue constitute a reliable methodology.

## DISCUSSIONS

This manuscript demonstrates direct measurement of E-field magnitude, using BSO crystal based EO probes, in mice brain *in-vivo*. We introduced a novel method for *in-vivo* measurement of E-field in biological tissue, enabling accurate dosimetry studies in exact experimental scenarios.

### Medium preparation for calibration process

To calibrate our EO probes for E-field measurements in mice brain, we prepared a mixture of brain tissue extracted from several animals. It is possible that the dielectric properties of the ex-vivo brain does not match the *in-vivo* brain. Yet, extracting the brain samples close to the experiment time and keeping them in saline at 4°C kept the tissue as fresh as possible and the physical properties of the resulting sample remains as close as possible to those of the brain (See dielectric measurement description in the Materials and Methods section). However, the results obtained were in good agreement with E-field values computed from the temperature control experiments (Fig. 3F) and potential errors in the calibration process, including medium preparation, did not induce an error beyond the already existing cross-animal measurement variabilities (which we also observed in our temperature control experiments). Development of novel probe structures with established physics-based analytical expressions for effects of surrounding medium on induced E-field inside the probe could remove the need for calibration in brain tissue. For example, the E-fields inside a spherical object immersed in a background medium can be expressed by Rayleigh’s theory (^21^). Such new generation of probes could be calibrated by prepared material samples for measurement in any medium, and be used in different medium, provided that the physical properties are known.

### Prominent Applications

Our experiments, demonstrating direct measurement of RF E-fields *in-vivo*, introduce a framework that can be used and further developed for different applications, such as: 1) non-invasive measurement of RF E-Field: metal-inclusive probed are invasive to the distribution of the filed they are immersed in. EO probes provide the possibility of non-invasive direct measurement of the RF E-field and our study demonstrate the feasibility of their *in-vivo* application which can be employed in the following applications: a) MRI and RF safety dosimetry in animal models and in biological tissue or phantoms: *in-vivo* measurements in animals can be used for direct and accurate safety measurements but also to examine and fine-tune simulations. b) Dosimetry in proximity of medical implants: one of the main safety concerns about RF applications (such as MRI, inspection devices, RF equipment, etc.) is the potential danger for people and patients with metallic implants. Animal models can be used to study such effects in live tissue rather than present approaches relying on *ex-vivo*, phantom or simulation studies. In this regard, using EO probe technology can be highly desirable given the fact that these probes minimally interact with the EM field. 2) low power dosimetry: Optical temperature measurements have also been shown to be a suitable way to provide an indirect estimate of the E-fields, but they application is reliable only for higher level of RF power where adequate temperature change is induced in the tissue. EO probes are the appropriate choice for measurements of the electric field induced by low power RF energy in the tissue.

### Uncertainties and Improvements

The results of our measurements demonstrate a noticeable extent of variability. We attribute this variability to various parameters including: 1) animal to animal difference: variability in age, body and head size, tissue (skin, fat, etc) variabilities between animals lead to an extended distribution of the induced E-field at the point of measurement. 2) Probe replacement: due to the three-step measurement procedure (Fig. 3B), the impact of the error of misplacement of the probe is increased. Development of three-axial probes can reduce this error. 3) Probe size versus mouse head volume: although probe dimensions in this set of experiments are small enough to allow implanting *in-vivo* and obtain relatively accurate measurements, the 1mm size of the probe adds some variability on the measurement from one animal to other. Development of sub-millimeter probes can help reduce this effect. 4) Dielectric and thermal property estimation: both experiments and simulations utilized information about the media in order to estimate the electric field (either by calibrating in phantoms with known thermal and dielectric properties, or via conduction of E-field simulations). A >10% uncertainty in the estimation of both the thermal and dielectric properties of the brain tissue has been reported in past studies (^22^) and is incumbent on this study as well. With these uncertainties in mind the E-field measurements exhibited an inter-animal variability of 16.3%. These errors are generally low, as for example in cellphone compliance testing conducted in phantoms, the maximum allotted errors are on the order of 30% (^23^).

While our experiments successfully demonstrate the possibility of direct measurement of RF E-fields *in-vivo*, further developments for improving the E-Field probes can ameliorate the quality and accuracy of the measurement. Some of such potential improvements include: 1) the development of a 3-axis probe that would reduce variability in (re)implantation and measurement procedure; 2) smaller probe footprint, enabling smaller interaction for reducing of field perturbation leading more accurate E-field readings with a decrease in tissue damage; 3) integration of EO E-field and optical temperature measurement: such technology would allow self-calibration of E-field values with temperature change measured at the same location in space.

Future extension of this work can be used for a more direct and accurate measurement of RF E-field in higher frequencies such as millimeter-wave, where RF energy is absorbed in shallow depths (given the skin-effect) and studies into in-vivo dosimetry is interest of the research field.

## MATERIALS AND METHODS

### Experiments

#### Phantom material preparation

We mixed the following materials in mass proportion until a homogenous solution is obtained: 130 gr (or 0.64% mass ratio) Corn Syrup (Golden, King Syrup, USA), 70 gr (or 0.35% mass ratio) Distilled water, 2 gr (or 1% mass ratio) salt (NaCl). The density (ρ), relative permittivity (ε), electrical conductivity (σ) and heat capacitance (C) were measured giving 1233.23 (Kg/m^3^), 47.84 (a.u.) at 950MHz, 1.06 (S/m) at 950MHz and 4514.17 (J/Kg.°C), respectively. Relative permittivity and electrical conductivity were measured using a dielectric probe kit (85070 E, Agilent Tech. Inc.) for a range of frequency between 950 MHz and 2400 MHz and heat capacitance was measured using a KD2 thermal properties analyzer (Decagon Devices Inc.).

#### Brain tissue preparation

We collected 219 mice, different types both male and female, from various laboratories in our institution, that were on the waiting list to be sacrificed, mostly because they were untagged animals from breeding of genetically modified animal lines. This approach avoided purchasing and killing extra animals. Mice were kept in cages in a 12hr regular cycle vivarium room dedicated to mice in up to five-occupancy cages. No prior experimentation had been performed on the animals. Mice were sacrificed the day before the probe calibration experiment and their brains were extracted. Brain were kept in saline at 4°C overnight. Prior to the experiment saline is extracted and brain tissue were mixed. We measured the following physical properties of the brain tissue mixture: density (ρ), relative permittivity (ε), electrical conductivity (σ) and heat capacitance (C) and obtained 988.93 (Kg/m^3^), 57.77 (a.u.) at 950Mhz, 1.06 (S/m) at 950MHz and 3540 (J/Kg.°C), respectively. Relative permittivity and electrical conductivity were measured using a dielectric probe kit (85070 E, Agilent Tech. Inc.) for a range of frequency between 950 MHz and 2400 MHz and heat capacitance was measured using a KD2 thermal properties analyzer (Decagon Devices Inc.).

#### Sensor calibration for E-field measurement in a medium

For calibration of the EO E-field probes, we positioned them at the center of a reservoir filled with the medium of interest and placed in the middle of one of the two chambers of a TEM cell antenna (TBTC0, Tekbox Digital Solutions). A specific set-up configuration was used for calibration of tangential or normal probes as shown in Fig. 2C. The TEM cell was fed by RF energy generated by a signal generator (845-26, BNC Berkeley Nucleonic) and an amplifier (ZHL-100W-13+, Mini-circuits). After setting-up the experiment configuration RF energy was injected into the TEM cell (36W input power) and recorded the uncalibrated E-field value using the E-field measurement system (NeoScan, EMAG Tech. Inc) for at least 20s (1Hz sample rate). The E-field probe was then removed and replaced by an optical temperature probe (0.3mm diameter, PRB-329 100-01M-STM, Osensa Innovation Corp.) in the same position, where temperature was recorded for 50s of baseline (no RF energy applied) followed by 70s of RF energy application into the TEM cell (36W input power). RF power was monitored using a power meter (U2001A Power Sensor and N9912A Field Fox Spectrum analyzer, Agilent/Keysight) that measured the forward and reflected power on a bidirectional coupler (778D, Agilent/Keysight) that was in line with the transmit chain.

#### Numerical simulations

A 3D implementation of our probe calibration set-up, consisting of a TEM cell antenna with a reservoir filled with a medium of interest placed in the middle of one of its chambers, was created in COMSOL Multiphysics (Comsol Inc.). A 3D EO probe, consisting of a 1 mm^3^ cubic BSO crystal attached to a cylindrical glass optic-fiber, was placed at the same location as in our probe calibration experiment for the normal probe. We defined virtual probing volumes at the center of the BSO crystal and adjacent to its bottom and lateral walls to monitor the E-field (measured as average over the volume) at different locations. First, simulations were conducted with a parametric sweep on the input voltage (and hence the input power) of the TEM cell. Then, at a fixed input power to the TEM cell (36W as in our probe calibration experiments) we performed a 2D parametric sweep, where the relative permittivity was changed from 1 to 78.5 (a.u.) with 2.5 (a.u.) increments and the electrical conductivity was changed from 0.4 to 4 (S.m^-1^) with 0.2 (S.m^-1^) increments, to evaluate the effects of the dielectric properties of the medium on the E-field sensing of the EO probe. Results of E-field virtual probing volumes were plotted following completion of the simulations.

#### Animals for the in-vivo E-field measurements

Adult wild-type male C57BL/6JxFVB mice (20-24 gr) were obtained from Charles River Laboratory. Mice were kept in cages in a 12hr regular cycle vivarium room dedicated to mice in up to five-occupancy cages. In all experiments, each animal served as its own control, no randomization or blinding was employed. No prior experimentation had been performed on the animals. All experiments were conducted in accordance with the Institutional Animal Care and Use Committee (IACUC) of New York University Medical Center.

#### EO E-field probe implantation and In-vivo RF E-field measurement

The implantation surgery was performed in two separate steps. Two days before the experiment, mice were anesthetized by isoflurane and a head-cap base matching the head-fixing set-up was attached to their skulls. In the day of the experiment, mice were anesthetized by urethane (1.5 g/kg, intraperitoneal injection) and positioned on the head-fixing set-up next to a stereotaxic system. A craniotomy was made and using the stereotaxic system the E Field probes were located inside the brain at a fixed location (similar to the location of the imaging lens in a former related experiment (^20^), at AP −2.1 mm, ML 1.5 mm, DV 1.5 mm coordinates from the Bregma point of the skull). The probes were secured by a custom-made 3D printed holder fixed to the head-fixing set-up (Fig. 3A). The set-up was then freed from the stereotaxic system and was moved to the adjacent room where the RF stimulation was performed. The patch antenna was placed 4cm away from the animal’s head (Fig. 3A) in the exact way as used in our former study (^20^). E-field measurement was done by two types of E-field probes: 1) normal probe which measures the E-field along the central axis of the probe and 2) tangential probe which measures the E-field along a certain direction perpendicular to the axis of the probe. To measure three orthogonal components of the E-field inside the brain the following process was used: First the tangential probe was implanted and the E Field along a direction perpendicular to the probes axis is measured (E_X_). Then, the probe was taken out and turned 90° and E Field in a perpendicular direction relative to the first direction and to the axis of the probe was measured (E_Y_). Finally, the tangential probe was taken out and the normal probe was implanted and the E-field along the axis of the probe was measured (E_Z_). Note that E_X_, E_Y_, and E_Z_ are the measured values after applying their respective calibration factors for the corresponding E-field probes. The total E-field is expressed as 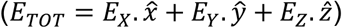 with a magnitude computed according 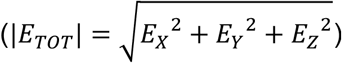. Animals are humanely euthanized at the end of the experiment.

### Analysis

#### Calculation SAR and E-Field from Temperature Measurements

Temperature changes induced by continuous-wave RF exposure (Fig. 2F) was used to calculate SAR and E-field values using 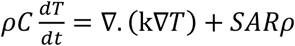 (Eq. 4), where ρ is the tissue density (kg/m3), C is heat capacity (J/kg/C), k is thermal conductivity (W/m/C), and SAR (W/kg) is the driving force for temperature rise defined as: *SAR* = *σ*|*E*|^2^/2*ρ* (Eq. 5). Where E is the induced E-field (V/m), and σ is the electrical conductivity (S/m). Under the short-term heating regime, Eq. 4 is simplified to: *SAR* = *C*Δ*T*/Δ*t* (Eq. 6) where ΔT/Δt is the temporal rate of temperature increase. Therefore, knowledge of the dielectric properties and thermal properties of the tissue, and heating duration (measured as specified above), as well as the temperature changes can effectively be used to estimate the magnitude of the electric field, provided it is dominated by a single polarization. SAR was calculated from the temporal temperature increase rate using Eq. 6 and E-field was driven from SAR values using Eq. 5 (^19^).

## Acknowledgements

Authors thank Mihaly Voroslakos for his help in the early establishment of this collaboration. O.Y. thanks Chiung-Yin Chung, Jingjing Liu, Pavan Veeramreddy, Riccardo Melani, Gergely Komlosi and Marisol Soula for their help with brain tissue preparation. This work was supported by the following grants: NIH-R01 (# 1R01NS113782-01A1) and NIH-TL1 postdoctoral fellowship (#2TL1TR001447-06A1) to O.Y.

## Author contributions

O.Y. and G.B. conceived the project. O.Y. designed the experiments. O.Y. prepared the animal brain samples and performed dielectric property measurements. F.S and O.Y. prepared the phantom material and performed dielectric property measurements. O.Y. designed and performed all animal experiments with help from L.A. and A.Sar. S.S., A.Sar. and A.Sab. conducted E-field probe preparation and E-field measurements. O.Y. performed all temperature measurements. O.Y. analyzed the data for all experiments. L.A. and O.Y. designed and performed the numerical simulations. O.Y. and K.S. wrote the paper and all authors participated in its revision. O.Y., G.B., L.A., and F.S. declare no competing interest. S.S., A.Sar., A.Sab., and K.S are working with EMAG Technologies Inc.

## SUPPLEMENTARY INFORMATION

**Supplementary Figure 1.**
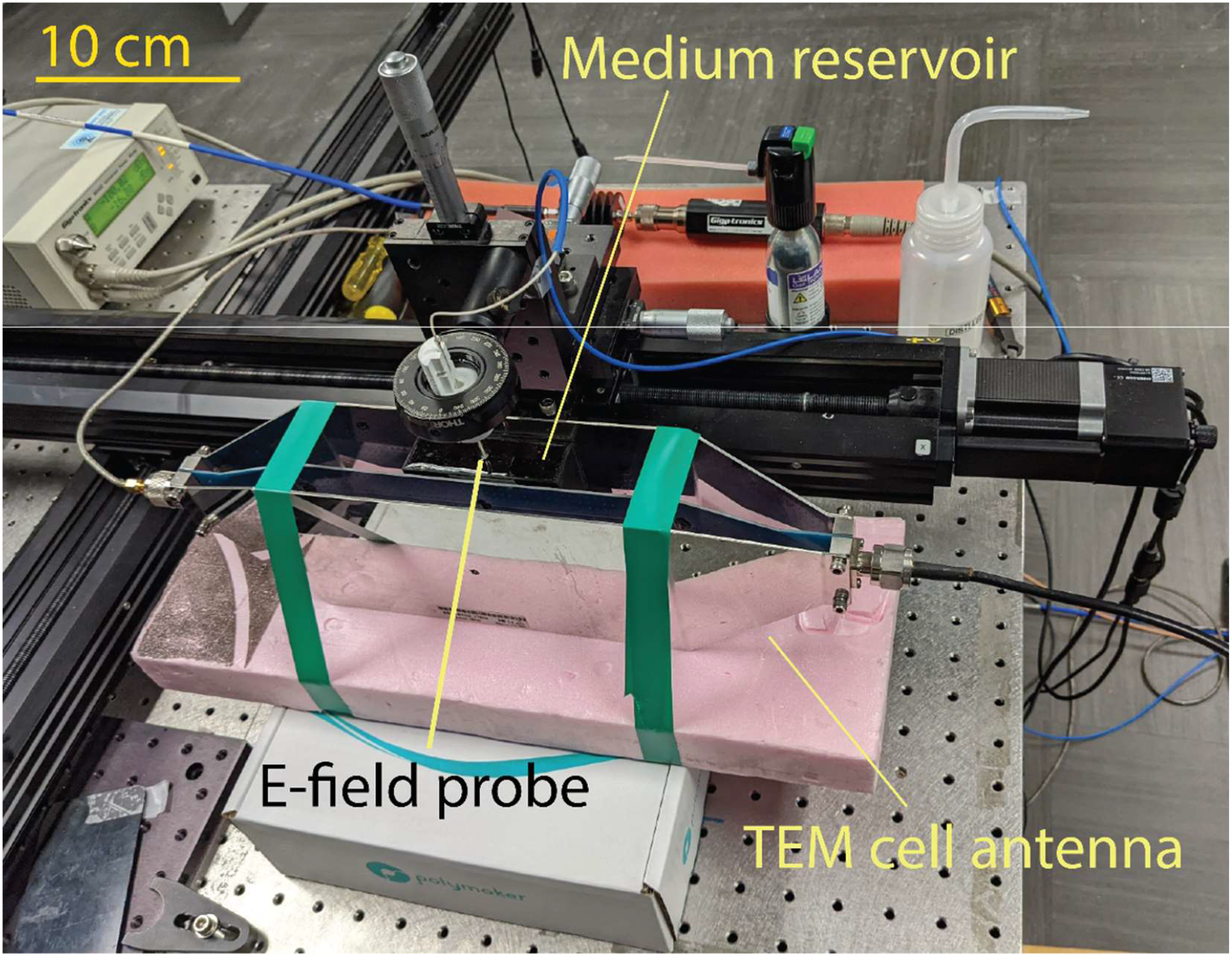
Probe calibration setup. Photo showing the calibration set-up configuration for the tangential probe using a TEM cell antenna. Here, the medium reservoir is filled with phantom material preparation.

**Supplementary Figure 2.**
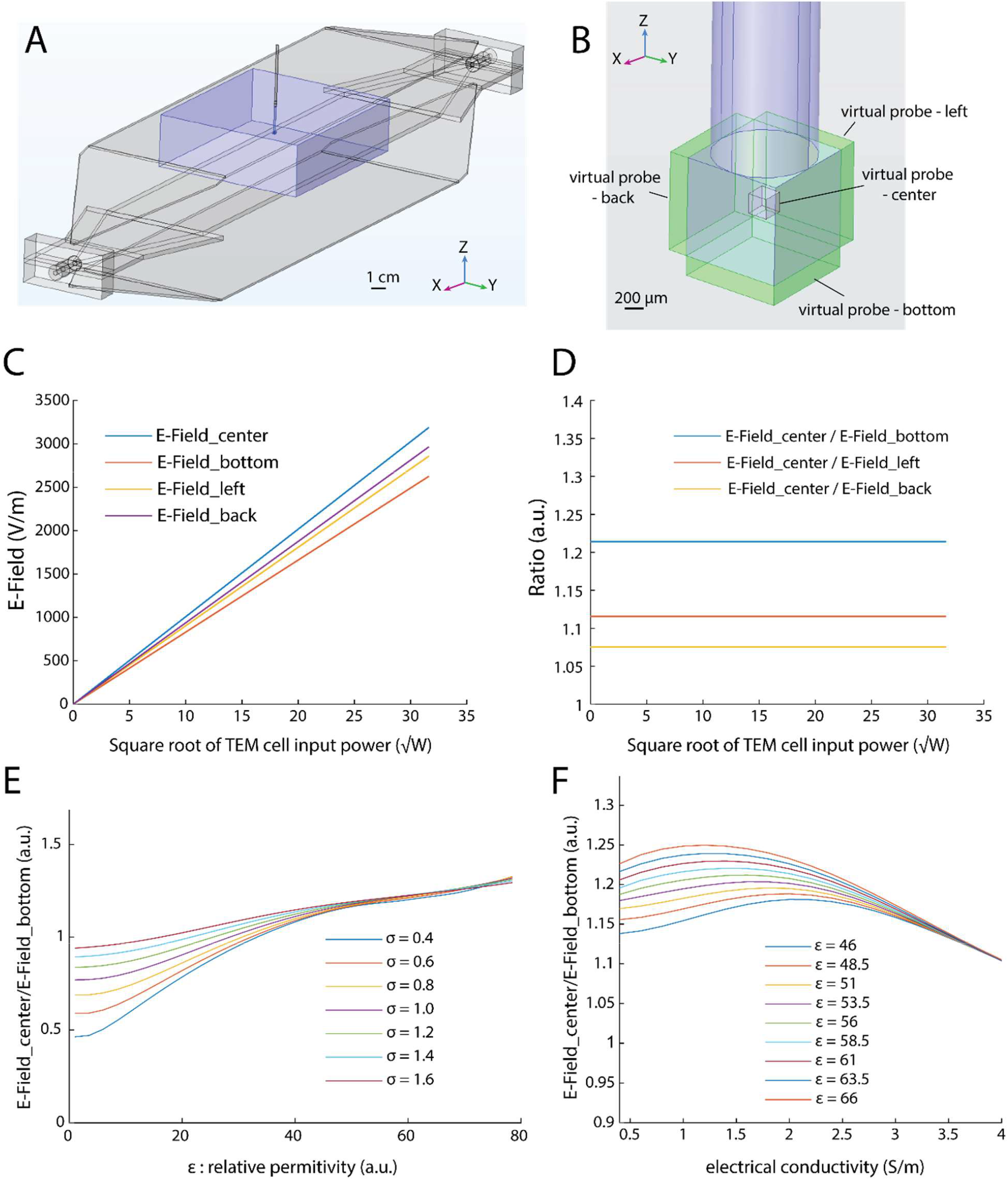
Numerical simulations. **A)** 3D drawing of the simulation set-up mimicking the calibration process for the normal probe including a TEM cell with the reservoir of the medium of interest (in blue). **B)** Zoomed-in view of the inserted probe (as in A) with the defined E-Field measurement probing volumes at the center and adjacent to the BSO crystal. Note that here ‘virtual probe’ regions (green) represent simulation tools to measure the average value of a parameter (here the E-field) across a defined volume. **C and D)** Measured electric-filed values for all probing volumes show a linear relationship with the square root of the input power to the TEM cell (using the electrical conductivity of 1.06 (S/m) and relative permittivity of 57.77 (a.u.)). **E)** Variation of the ratio between the E-field at the center of the BSO crystal and the in the probing volume beneath it (this is the area that the normal probe is supposed to monitor) as a function of the relative permittivity of the medium of interest. Note the relatively low variability of this ratio for different electrical conductivity of the medium at relative permittivity values around that of the brain tissue (∼57). **F)** Same measure as in E but as a function of the electrical conductivity of the medium of interest. Note the relatively low variability of this ratio for different relative permittivity of the medium at electrical conductivity values around that of the brain tissue (1 S/m).

**Supplementary Figure 3.**
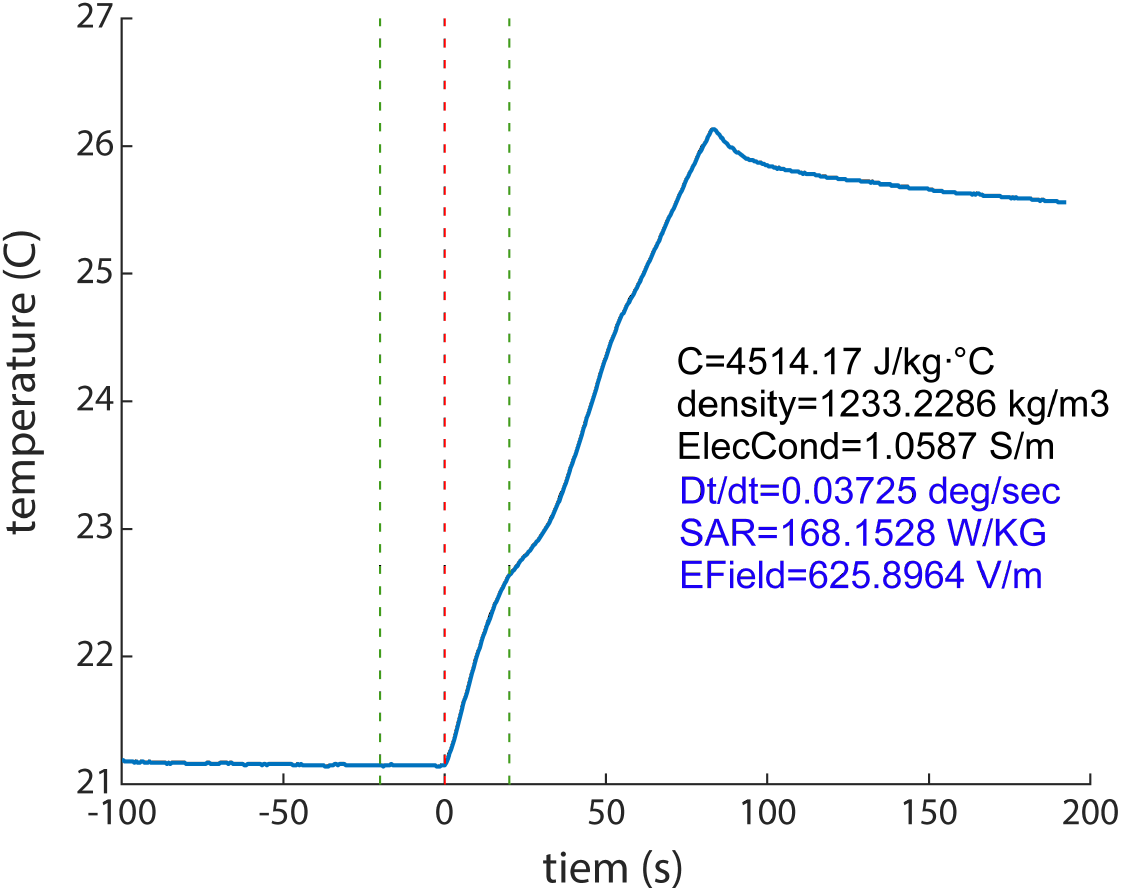
Temperature measurement for probe calibration in Phantom preparation. Example temperature measurement for E-Field estimate calculation for calibration of E-Field probes immersed in a reservoir filled with a brain phantom preparation in a TEM cell.

**Supplementary Figure 4.**
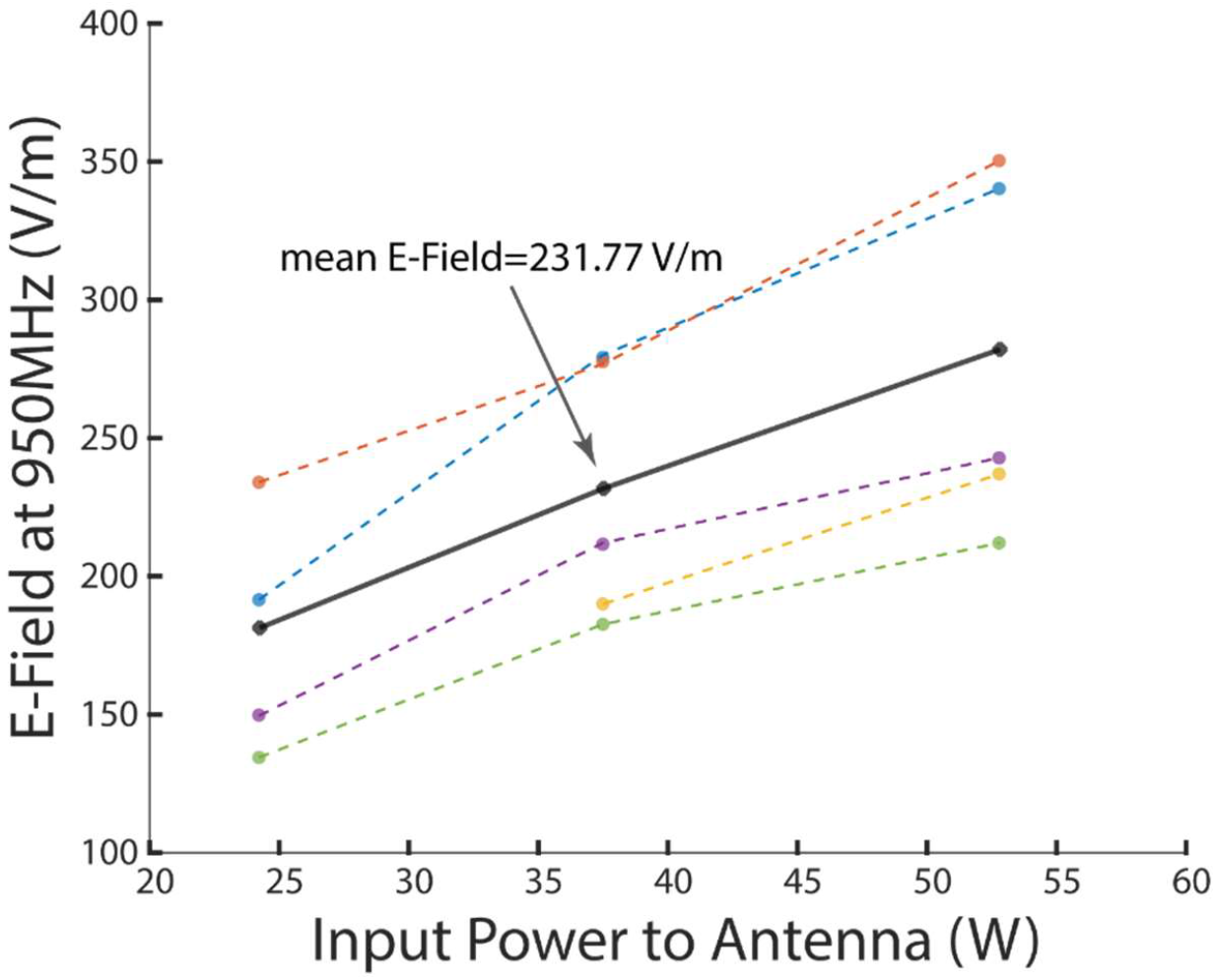
E-field estimate from in-vivo temperature measurements. Data from (Yaghmazadeh2022).

## REFERENCES

1. Daily, L. E. A clinical study of the results of exposure of laboratory personnel to radar and high frequency radio. Nav. Med. Bull. 41, 1052 (1943).

2. Follis, R. H. Studies on the biological effect of high frequency radio waves (radar). Am. J. Physiol. 147, 281 (1946).

3. Michaelson, S. M. & Lin, J. C. Biological Effects and Health Implications of Radiofrequency Radiation. (Plenum Press, 1987).

4. Vöröslakos, M., Yaghmazadeh, O., Alon, L., Sodickson, D. K. & Buzsáki, G. Brain-implanted conductors amplify radiofrequency fields in rodents: advantages and risks. bioRxiv (2022).

5. Karipidis, K., Mate, R., Urban, D., Tinker, R. & Wood, A. 5G mobile networks and health—a state-of-the-science review of the research into low-level RF fields above 6 GHz. J. Expo. Sci. Environ. Epidemiol. 31, 585–605 (2021).

6. Dagro, A. M., Wilkerson, J. W., Thomas, T. P., Kalinosky, B. T. & Payne, J. A. Computational modeling investigation of pulsed high peak power microwaves and the potential for traumatic brain injury. Sci. Adv. 7, 1–11 (2021).

7. Foster, K. R. Thermal and nonthermal mechanisms of interaction of radio-frequency energy with biological systems. IEEE Trans. Plasma Sci. 28, 15–23 (2000).

8. Ziegelberger, G. et al. Guidelines for limiting exposure to electromagnetic fields (100 kHz to 300 GHz). Health Physics vol. 118 (2020).

9. Gabriel, C. & Gabriel, S. Compilation of the Dielectric Properties of Body Tissues at RF and Microwave Frequencies. (1996).

10. Gabriel, C., Gabriel, S. & Corthout, E. The dielectric properties of biological tissues: I. Literature survey. Phys. Med. Biol. 41, 2231–2249 (1996).

11. Schuderer, J., Schmid, T., Urban, G., Samaras, T. & Kuster, N. Novel high-resolution temperature probe for radiofrequency dosimetry. Phys. Med. Biol. 49, (2004).

12. Faraone, A., McCoy, D. O., Chou, C. K. & Balzano, Q. Characterization of miniaturized E-field probes for SAR measurements. IEEE Int. Symp. Electromagn. Compat. 2, 749–754 (2000).

13. Cecelja, F. & Balachandran, W. Electrooptic Sensor for Near-Field Measurement. IEEE Trans. Instrum. Meas. 48, 650–653 (1999).

14. Wu Q. Z. X.-C. 7 terahertz broadband GaP electro-optic sensor. Appl. Phys. Lett. 70, 1784–1786 (1997).

15. Keiber, S. et al. Electro-optic sampling of near-infrared waveforms. Nat. Photonics 10, 159–162 (2016).

16. Sulzer, P. et al. Determination of the electric field and its Hilbert transform in femtosecond electro-optic sampling. Phys. Rev. A 101, 1–17 (2020).

17. Sarabandi, K., Choi, J., Sabet, A. & Sabet, K. Pattern and Gain Characterization Using Nonintrusive Very-Near-Field Electro-Optical Measurements over Arbitrary Closed Surfaces. IEEE Trans. Antennas Propag. 65, 489–497 (2017).

18. Jarrige, P. et al. Electrooptic probe adapted for bioelectromagnetic experimental investigations. IEEE Trans. Instrum. Meas. 61, 2051–2058 (2012).

19. Alon, L., Sodickson, D. K. & Deniz, C. M. Heat Equation Inversion Framework for Average SAR Calculation From Magnetic Resonance Thermal Imaging. Biolectromagnetics 176, 139–148 (2016).

20. Yaghmazadeh, O. et al. Neuronal activity under transcranial radio-frequency stimulation in metal-free rodent brains in-vivo. Commun. Eng. 1, (2022).

21. Bohren, C. F. & Huffman, D. R. Absorption and scattering by a shpere. in Absorption and scattering of light by small particles 82–129 (WILEY-VCH Verlag GmbH & Co. KGaA, 1998).

22. Sasaki, K., Porter, E., Rashed, E. A., Farrugia, L. & Schmid, G. Measurement and image-based estimation of dielectric properties of biological tissues —past, present, and future—. Phys. Med. Biol. (2022) doi:10.1088/1361-6560/ac7b64.

23. Marinescu, A. Measurement Uncertainty at SAR Determination for Mobile Phones. in The 6th International Symposium on ADVANCED TOPICS IN ELECTRICAL ENGINEERING 1–5 (2008).

